# Environmental microbial communities and host selection shape larval microbiomes

**DOI:** 10.64898/2026.05.14.725214

**Authors:** Stephanie F. Hendricks, Amy L. Tan, Amelia G. Williams, Katherine M. Buckley, Marie E. Strader

## Abstract

Ocean warming is altering abiotic environments and biotic interactions experienced by marine organisms, where sensitive early developmental windows occur in biologically complex seawater communities. The impact of these interactions on developmental processes and fitness in hosts is not well understood, but likely contingent on the establishment of a host-associated microbiome. Here, we hypothesize that temperature and microbial exposure during embryogenesis influence larval microbiome assembly and host morphology. *Strongylocentrotus purpuratus* embryos were raised in low microbial richness (LMR) or high microbial richness (HMR) seawater at ambient (14 °C) or elevated (18 °C) temperature, then collected at 2, 4, and 6 days post-fertilization (dpf) following multiple feedings. Higher microbial diversity was observed in larvae that developed in HMR seawater when compared to LMR. Differences in relative abundances of dominant microbial families between seawater and larvae suggest some degree of host selectivity in microbiome assembly. Temperature did not strongly alter microbiome composition, but both temperature and microbial condition led to differences in larval morphology by 6 dpf, potentially due to enrichment of microbes with chemoheterotrophic functions. By linking how temperature and microbial communities interact with host development, we contribute novel insights into how early-life environmental conditions impact holobiont formation and morphology.

**One sentence summary:** Early developmental temperature and microbial conditions shape larval microbiome establishment and morphology.

## Introduction

The holobiont, defined as the host and its associated microbial communities (Bordenstein & Theis, 2015; Simon et al., 2019), plays a central role in mediating organismal responses to environmental changes. Environmental conditions experienced early in development may have long-lasting phenotypic effects (Burton & Metcalfe, 2014), and growing evidence suggests that internal and external microbial communities contribute to these effects (Fallet et al., 2022; Koide, 2023; Dantan et al., 2024; Pepke et al., 2024; Tan et al., 2025). Microbiomes can modulate many host traits by influencing host metabolism and growth, immune function, and developmental processes (McFall-Ngai et al., 2013; Schuh et al., 2020; Park et al., 2023). Meanwhile, both establishment of the microbiome and maturation of the holobiont are influenced by environmental factors (*e.g.,* temperature, salinity, habitat type), local microbial environments, and host-associated factors such as diet and genetics (Abdelfadil et al., 2024; Bengtsson et al., 2025; Bright & Bulgheresi, 2010; Grieneisen et al., 2021; Lima et al., 2020; Pantos et al., 2015). Thus, in the holobiont, the microbiome acts as an interface between the organism and its environment, which has profound implications for host resilience and adaptation, especially under conditions of rapid global change. However, it remains unclear how environmental variations and the formation of the microbiome interact to shape early life-history traits.

Early microbial colonization is often driven by environmental exposure, but hosts also influence microbial community composition through selective mechanisms, including immune responses and hormone signaling (Mazel et al., 2018). This selectivity results in microbial communities that are distinct from the surrounding environment (Angthong et al., 2020; Chen et al., 2024; Haditomo et al., 2021; Hakim et al., 2019; Rodríguez-Barreras et al., 2021; Sadeghi et al., 2023; Salonen et al., 2021). Microbiome selectivity, however, varies immensely across species and developmental stages (Koskella & Bergelson, 2020), ranging from lack of stable, resident microbiomes to transient selection that changes with life stage to highly specific interactions that are crucial for host functions (Engel & Moran, 2013; Hammer et al., 2017; Kwong & Moran, 2016; Marangon et al., 2023; Martinson et al., 2011; Muffett et al., 2025; Nyholm & McFall-Ngai, 2021). Understanding the interplay between environmental exposure and host selection that shapes the microbiome is essential for understanding how the holobiont assembles and functions.

Host-microbe interactions are especially important during early-life stages when organisms are particularly vulnerable to environmental stressors (Angthong et al., 2020; Fallet et al., 2022). In many marine invertebrates, microbial communities can induce phenotypic plasticity. Microbiome community composition and diversity are known to influence host development and life stage transitions (Cavalcanti et al., 2020), growth and metabolism (Carrier & Reitzel, 2019; Donia et al., 2011), reproduction (Comizzoli et al., 2021; Stouthamer et al., 1999; Téfit et al., 2023), behavior (Archie & Tung, 2015), and resistance to pathogens and other stressors (Baquiran et al., 2025; Chuang et al., 2024; Koch & Schmid-Hempel, 2011; Quigley et al., 2023). Geographic and habitat-specific differences in microbial communities shape the microbes available for host colonization (Miller et al., 2021), which may expand the ways that animals can adjust to their environment (Henry et al., 2021). Planktonic larval marine invertebrates are typically exposed to the surrounding seawater from the time of fertilization; the microbial communities within this seawater may influence the ability to adjust to varying ocean conditions. In sea urchins, larval growth patterns change based on the microbial richness and diversity of the surrounding seawater (Carrier et al., 2019; Tan et al., 2025). Although recent work has described how the internal microbiome of the larval purple sea urchin forms and changes over time (French et al., 2024), it remains unknown whether the microbial communities influence larval plasticity. Together, these studies highlight how the environment and associated microbial exposure impact developing organisms, motivating a closer examination of how seawater microbiomes influence plasticity in early life-history traits.

Temperature is a strong driver of ocean microbial communities (Sunagawa et al., 2015; Pomeroy & Wiebe, 2001), and marine environments are experiencing increasingly intense and frequent marine heatwaves (MHWs) (Oliver et al., 2018). MHWs also include fluctuations in autocorrelated abiotic factors, including pH and dissolved oxygen (Mogen et al., 2022). Together, these strongly influence biotic factors such as shifts of planktonic communities (Brodeur et al., 2019; Eglaine et al., 2025). Consequently, shifts in seawater microbial communities driven by warming have the potential to reshape host-associated microbiomes, leading to plastic or even permanent phenotypic variation (Carrier & Reitzel, 2018). Here, we investigated how temperature and environmental microbial richness influence larval microbiome establishment and whether these factors drive phenotypic plasticity during early development. We maintained *Strongylocentrotus purpuratus* embryos at ambient (14 °C) or elevated (18 °C) temperatures in either low microbial richness (LMR) or high microbial richness (HMR) artificial seawater and collected at 2, 4, and 6 days post-fertilization (dpf). We used 16S rRNA amplicon sequencing to profile the microbial communities present in the surrounding seawater and microbiome establishment and progression in larvae. Further, we quantified larval morphometrics and identified the enrichment of specific microbial metabolic functions that may be associated with growth. Understanding how environmental factors shape microbiome establishment and morphology during early development is critical for predicting organismal responses to global change.

## Materials and Methods

### Adult animal husbandry, spawning, and fertilization

Gravid purple sea urchins (*Strongylocentrotus purpuratus*) were obtained from Point Loma Marine Invertebrate Lab (Lakeside, CA), Marinus (Garden Grove, CA), and Monterey Abalone Company (Monterey, CA) and maintained in aquaria at 15 °C, 32 ppt salinity in artificial seawater (ASW; Instant Ocean) at Auburn University, Auburn, AL. Sea urchins were fed kelp (*Macrocystis pyrifera*) once per week. Spawning was induced by manual agitation or intracoelomic injection of 0.53 M KCl. Gametes were collected from three males and three females to create three unique genetic crosses, which represented biological triplicates throughout the experiment. Therefore, each cross was represented in every treatment condition. This design enabled us to confirm consistency of the treatment effect across multiple familial crosses, given that host genotype can strongly influence microbiome composition. Sperm was collected dry and placed on ice until activation; eggs were collected in 15 °C, 30 ppt salinity filter-sterilized (0.2 µm) ASW. Following test fertilizations to confirm > 95% fertilization success, three separate fertilizations were generated from the three unique pairs of animals. Fertilized eggs from each cross were evenly divided into four culture vessels (*i.e*., four treatment conditions).

### Embryo cultures

Embryos from each of the three genetic crosses were cultured at a density of ~10 embryos/mL in 12 L chambers (~120 000 total embryos in each chamber) at 30 ppt salinity. Cultures were maintained in temperature-controlled incubators at either ambient (14 °C) or elevated (18 °C) temperatures in ASW collected from the aquaria housing the adult sea urchins and treated in one of the following ways to adjust microbial content: 1) low microbial richness (LMR) seawater was filter-sterilized (0.2 µm) to remove most bacteria; or 2) high microbial richness (HMR) was mesh-filtered (40 µm) to remove large particles and debris but retain microbes (Figure 1A). To prevent contamination and bacterial growth, algae (*Rhodomonas lens*) were washed in filter-sterilized (0.2 µm) ASW. Larvae were fed algae (3000 cells/mL of larvae culture) on days 3 and 5. Water changes were performed (1/3^rd^ volume) using the appropriately treated seawater every other day.

**Figure 1.**
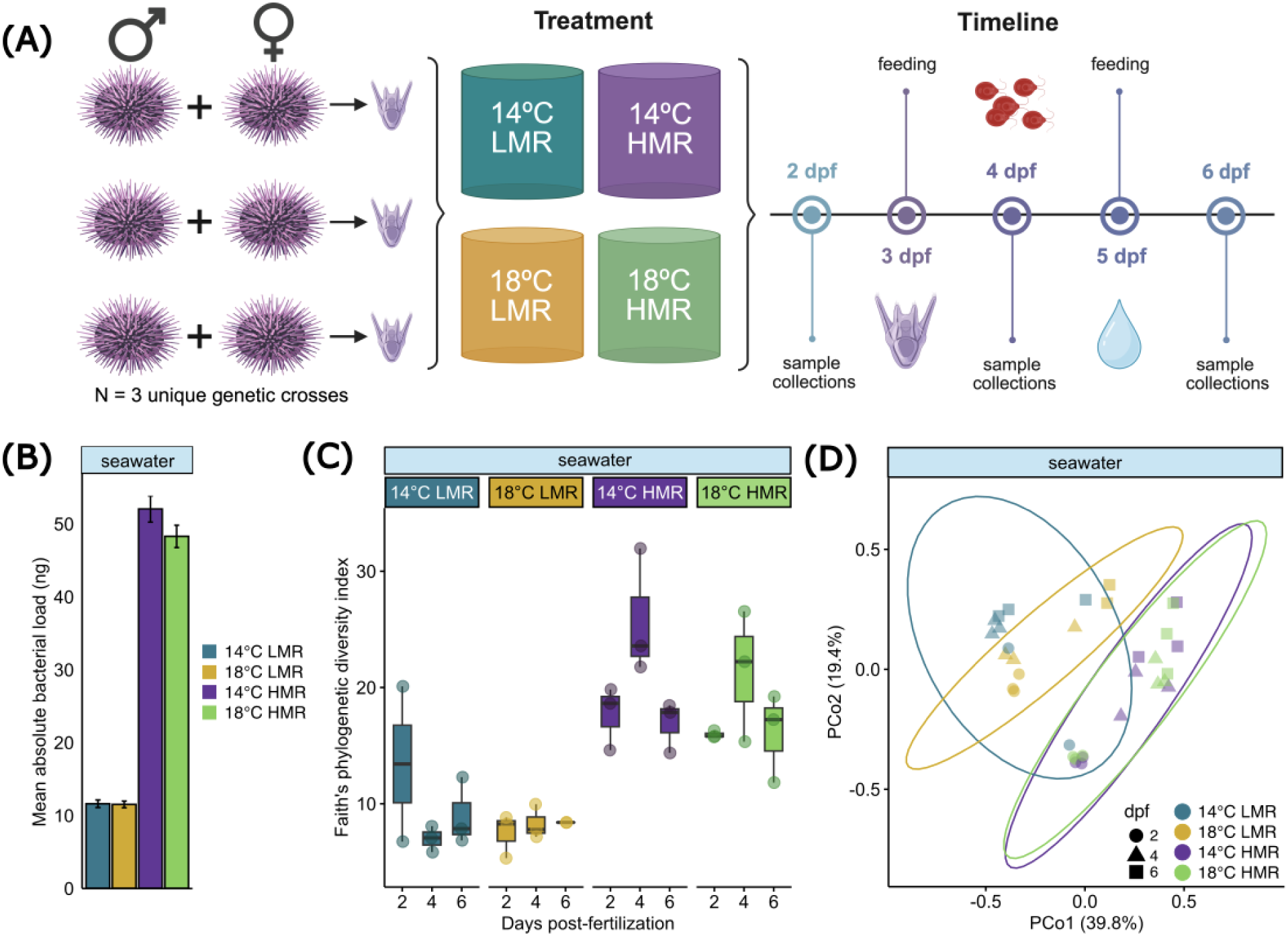
Seawater microbial treatments exhibit differences in microbial load and diversity. A) Embryos from three unique genetic crosses, which served as biological triplicates, were subjected to one of four treatments: two temperatures (14 °C or 18 °C) and two seawater conditions (low microbial richness [LMR] or high microbial richness [HMR]). Larvae and seawater samples were collected at 2, 4, and 6 days post-fertilization (dpf) for 16S rRNA sequencing to conduct microbial community analysis. Larvae were fed with *Rhodomonas lens* at 3 and 5 dpf. B) Mean absolute microbial load from qPCR across four treatments at 6 dpf in seawater samples. Error bars indicate standard deviation. C) Faith’s phylogenetic alpha diversity across four treatments at all time points in seawater samples. Each box represents the interquartile range (IQR) with the median line, whiskers extend to 1.5× the IQR. Each point is a sample. D) Bray-Curtis beta diversity across four treatments at all time points in seawater samples. Ellipses represent 95% confidence intervals. Each point is a sample and shape indicates time point.

### Sample collection

To characterize the larval-associated microbiome, larvae (n = 5000) were collected at 2, 4, and 6 days post-fertilization (dpf) and pelleted via centrifugation. All seawater was removed to limit contamination, and the pellets were immediately flash-frozen in liquid nitrogen. Collection equipment was rinsed with sterile seawater between collections from different larval chambers to avoid cross-contamination. To assess larval morphology, larvae (6 dpf; n = 200) were collected and preserved in Z-fix (Anatech). After each larval sample collection, 300 mL of seawater from each culture vessel was vacuum-filtered using a 0.45 µm filter paper in a Büchner funnel to capture environmental microbes.

### DNA extraction and quality control

Microbial DNA was extracted from the filters (n = 36) using a modified version of the ZymoBIOMICS DNA Miniprep Kit protocol with an extended digestion period (Tan et al., 2025). One seawater DNA sample was not sent for sequencing analysis due to low DNA concentration, and two samples were pooled together from the same environmental condition and time point due to low concentration (total of n = 34 samples sequenced). Larval DNA (n = 36) was extracted using a CTAB (Cetyltrimethylammonium bromide) protocol (Murray & Thompson, 1980; Saghai-Maroof et al., 1984). Two larval DNA samples were pooled together from the same environmental condition and time point before sequencing due to low concentration (total of n = 35 samples sequenced). Isolated DNA was quantified using a Qubit fluorometer.

### Quantification of bacterial load

Bacterial load of the seawater samples was quantified using qPCR. Due to the overall low concentrations of DNA from the seawater, samples were pooled based on treatment condition (*e.g.*, equal volume of the DNA extractions from the three 14 °C LMR larval chambers). The TaqMan 16S pan-bacterial assay (Ba0490791_s1, ThermoFisher) was run on a BioRad CFX Opus 96 along with a standard curve created by serial dilutions of the ZymoBIOMICS Microbial Community DNA. Standard. C_t_ values of the seawater samples and standard curve were compared to calculate absolute bacterial load. Assumptions of normality and homogeneity were tested using Shapiro-Wilk and Levene tests using packages *stats* (R Core Team, 2023) and *car* (Fox & Weisberg, 2019), respectively. As a result of these tests, an aligned rank transform (ART) (function “art”) was run to prepare for a nonparametric analysis of variance (ANOVA) with formula temperature × seawater condition, followed by a contrast test (Elkin et al., 2021; Kay et al., 2025; Wobbrock et al., 2011).

### 16S amplicon sequencing

Microbial communities within the seawater and larvae were assessed using 16S amplicon sequencing on the isolated DNA. DNA samples were sent to Novogene for barcoding, amplification, and library preparation as described previously (Tan et al., 2025). Amplicons were sequenced on the NovaSeq 6000 platform (250-bp paired-end reads). On average, 197 998 raw paired-end reads were generated across all samples.

### Larval morphology analysis

Z-stack brightfield images of fixed larvae were taken on a wide-field Zeiss Axio Observer Fluorescent Microscope with Zen Blue imaging software and analyzed using ImageJ/FIJI (v1.53t) (Schindelin et al., 2012). To calculate larval length, the position of the apical point and the top of all four arms were marked. The x, y, and z position information of each point provided by the “Cell Counter” plugin was used to calculate the length of each arm based on the distance formula. The average of the four apex-to-arm measurements was used as the larval length. Three Z-stack images (BF18_0hpi_20x_stack9, TS18_0hpi_20x_stack1, TS18_0hpi_20x_stack4) were removed as outliers based on treatment. An interactive ANOVA was run with formula temperature × seawater condition, followed by a Tukey’s post hoc test.

### 16S analysis

To assess the microbial communities present in seawater and larval samples, 16S rRNA sequences (V3-V4 region) were analyzed using Quantitative Insights Into Microbial Ecology (QIIME2; v2024.10, (Bolyen et al., 2019), using SILVA 138.1 reference database with RESCRIPt (Pruesse et al., 2007; Quast et al., 2012; Robeson et al., 2021) specific to the V3-V4 region (see Tan et al., 2025 for more details). Two larval samples, BS14_6dpf_L and TS14_2dpf_L, were removed from analysis based on contamination from a single taxon (sequences assigned as *Xanthomonadaceae* accounted for >85% in these samples). In RStudio (v4.3.1), sequences identified as eukaryotic, chloroplast, or mitochondrial were removed. First, data were made into a “phyloseq” object with the function “qza_to_phyloseq” in package *qiime2R* (Bisanz, 2018). Using the package *phyloseq* (McMurdie & Holmes, 2013), samples were rarefied to the minimum sample depth of the lowest sequenced sample (82 246) without replacement using the function “rarefy_even_depth.” Relative abundances were calculated using the function “transform_sample_counts.” Alpha diversity was estimated with Faith’s phylogenetic diversity using the function “estimate_pd” in the package *btools* (Battaglia, 2018). To confirm that genotype did not have an impact on larval microbiomes, a Kruskal-Wallis test was run. A linear mixed effects model using the function “lmer” in package *lmerTest* (Kuznetsova et al., 2017) with formula temperature × water condition × day and chamber by day as a random effect was run. Significance of models was calculated with Satterthwaite’s method using the function “anova.” Model assumptions were assessed using simulated residual diagnostics implemented in the package *DHARMa* (Hartig et al., 2024), as well as visually inspected. Beta diversity was visualized with a principal coordinates analysis (PCoA) using Bray-Curtis with *phyloseq* functions “distance” and “ordinate,” and significance was tested using a PERMANOVA (package *vegan*, function “adonis2”) with 999 permutations (Anderson, 2001; McArdle & Anderson, 2001)). Package *ANCOM-BC* (Lin & Peddada, 2020) was implemented to analyze the differential abundance of taxa at the genus level in temperature (14 °C vs. 18 °C) and microbial seawater conditions (LMR vs. HMR), using all default parameters. In addition, to investigate the functional roles of microbes present in the seawater and larval samples across treatments, FAPROTAX (v1.2.11) was employed (Louca et al., 2016).

## Results

### Microbial communities vary significantly between seawater conditions

To ensure that larvae cultured in the two seawater conditions were exposed to different microbial abundances, we characterized the microbial load of the seawater from all four treatments using qPCR (Figure 1A). Results indicate that the HMR seawater contained significantly more microbial DNA than LMR (*p_ART ANOVA_* = 0.001; HMR was ~5X higher than LMR; Figure 1B). Temperature also had a significant effect on total microbial load (*p_ART ANOVA_* = 0.017; Figure 1B); however, temperature did not significantly alter microbial load within either seawater conditions (LMR and HMR).

To determine if this difference in microbial load also reflected a difference in the microbes present in the seawater, we amplified and sequenced the 16S rRNA gene for seawater collected at each of the three time points and four treatments. Temperature did not significantly alter microbial diversity in the seawater (*p* = 0.120) (Supplemental figure 1) nor did day (*i.e*., 2, 4, and 6 dpf; *p* = 0.290). However, microbial richness of the seawater condition was significant (*i.e*., LMR and HMR; *p* = < 0.001). When considering interactions, microbial richness and day were significant (seawater condition × day, *p* = 0.01). LMR seawater (14 °C LMR and 18 °C LMR) exhibited relatively stable, but low, alpha diversity across all time points (Figure 1C). In contrast, HMR seawater (14 °C HMR and 18 °C HMR) exhibited consistently higher alpha diversity than LMR (Figure 1C). Additionally, beta diversity differed significantly by microbial richness (*i.e.*, LMR and HMR; R^2^ = 0.29, *p*_PERMANOVA_ = 0.001) (Figure 1D). However, temperature (R^2^ = 0.03, *p*_PERMANOVA_ = 0.253) and the interaction of temperature and microbial richness (R^2^ = 0.04, *p*_PERMANOVA_ = 0.133) were not significant (Figure 1D). Taken together, seawater microbial community composition and diversity differed by microbial richness, but not temperature.

The microbial communities present within seawater were primarily composed of bacteria from nine families (Supplemental figure 2). Overall, the most abundant families represented were *Rhodobacteraceae*, *Vibrionaceae*, and *Flavobacteriaceae* (Supplemental figure 2). *Rhodobacteraceae* was present in high relative abundance in all treatments and time points (31% overall mean relative abundance; Supplemental figure 2). *Vibrionaceae* also represented a substantial proportion of the seawater microbial communities (14% overall mean relative abundance), with the exception of 18 °C LMR, in which it was relatively lower (Supplemental figure 2). *Pseudoalteromonadaceae* and *Alteromonadaceae* were also consistently present in all four treatments, but at lower relative abundances (12% and 7% overall mean relative abundance, respectively; Supplemental figure 2). In contrast, other families varied in abundance over time. For example, *Psychromonadaceae* was relatively rare on day 4 (0.4% overall mean relative abundance), and not present in 14 °C and 18 °C LMR on day 6 (Supplemental figure 2). *Flavobacteriaceae* was detected in the seawater on days 2 and 4 (0.9% and 13% overall mean relative abundance, respectively), but considerably increased by day 6 (33% overall mean relative abundance; Supplemental figure 2). Similarly, *Crocinitomicaceae* appeared by day 4 in all treatments except 14 °C LMR (< 0.06% overall mean relative abundance; Supplemental figure 2). In LMR conditions, *Shewanellaceae* and *Saccharospirillaceae* had greater relative abundances compared to HMR conditions (LMR was ~30X and ~8X higher, respectively; Supplemental figure 2). In spite of small shifts in the relative abundance of microbial families in the seawater treatments, the communities remained relatively consistent over time and were enriched primarily with microbial taxa found in natural seawater and associated with echinoderms (Bengtsson et al., 2025; Park et al., 2023; Yao et al., 2019; Yu et al., 2023).

### Larval microbiomes shift over time and are influenced by environmental conditions

To examine the establishment of the larval microbiome from early development through the onset of feeding, DNA was isolated from larval samples collected at 2, 4, and 6 dpf and used as the template to amplify and sequence the 16S rRNA gene. There was no significant difference in alpha diversity when testing for temperature (*i.e.*, 14 °C and 18 °C; *p* = 0.910) or day (*i.e.*, 2, 4, and 6 dpf; *p* = 0.510), however, seawater condition was significant (*i.e*., LMR and HMR; *p* = < 0.001). Further, alpha diversity was not significantly different for the interaction of temperature, microbial richness, and day (temperature × seawater condition × day, *p* = 0.590). Consistent with the seawater, microbiomes of larvae grown in LMR conditions (14 °C LMR and 18 °C LMR) displayed low but consistent alpha diversity across all time points (Figure 2A). Larval microbiomes from HMR conditions (14 °C HMR and 18 °C HMR) exhibited consistently higher alpha diversity over time, with no differences between triplicate chambers (Figure 2A, Supplemental figure 1F). This indicates that genotype did not significantly affect the larval microbiome (*p*_Kruskal-Wallis_ = 0.54). When considering beta diversity, microbial richness (R^2^ = 0.36, *p*_PERMANOVA_ = 0.001) and the interaction of temperature and microbial richness (R^2^ = 0.06, *p*_PERMANOVA_ = 0.018) were significant, but temperature alone was not (R^2^ = 0.03, *p*_PERMANOVA_ = 0.107) (Figure 2C).

**Figure 2.**
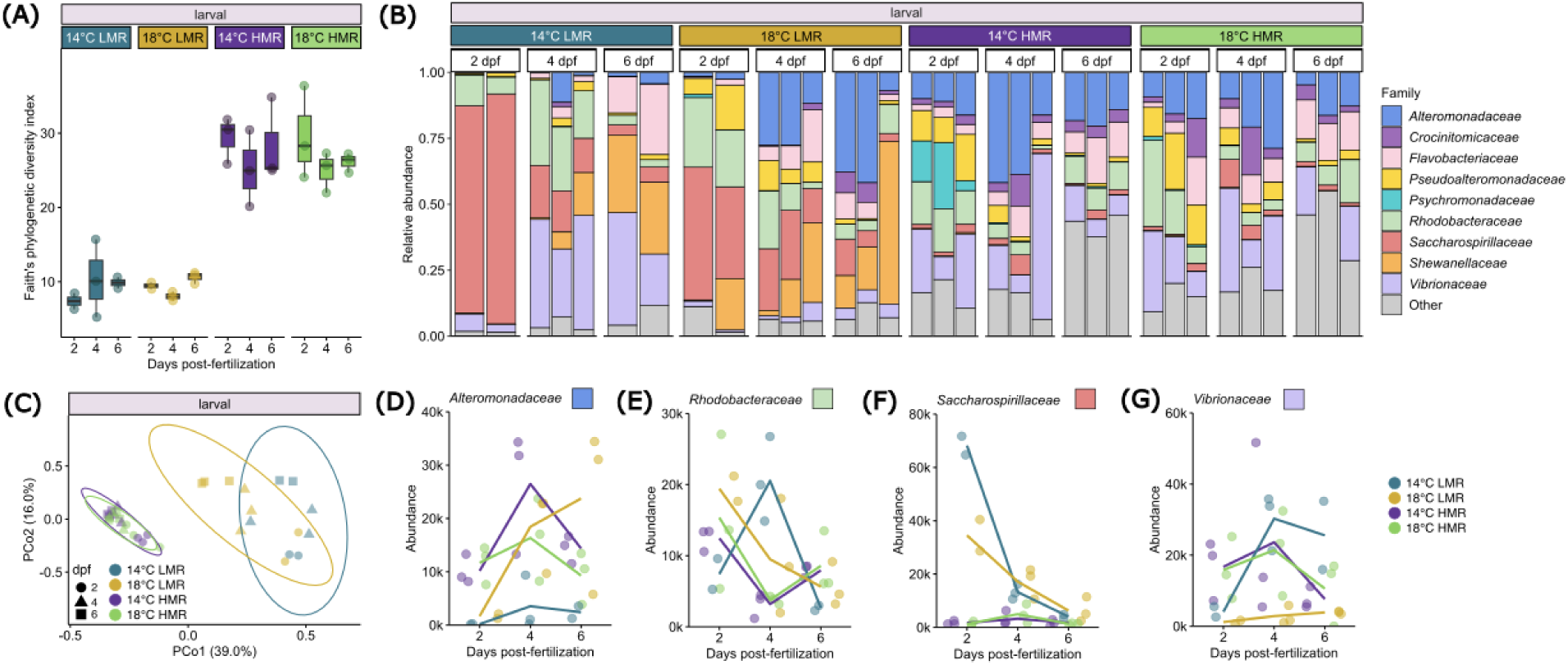
Larval microbiomes shift depending on environmental microbes and day. A) Faith’s phylogenetic alpha diversity across four treatments at all time points in larval samples. Each box represents the interquartile range (IQR) with the median line, whiskers extend to 1.5× the IQR. Each point is a sample. B) Stacked barplot with relative abundance of microbial communities across four treatments at all time points in larval samples. Each bar represents a sample, with colors indicating the top nine families (≥ 25% relative abundance in any single sample). “Other” consists of the remaining families grouped as one category. C) Bray-Curtis beta diversity across four treatments at all time points in larval samples. Ellipses represent 95% confidence intervals. Each point is a sample and shape indicates time point. (D-G) Line plots with abundance of *Alteromonadaceae* (D), *Rhodobacteraceae* (E), *Saccharospirillaceae* (F), and *Vibrionaceae* (G) across four treatments at all time points in larval samples. Each point is a sample.

Overall, the microbial families most commonly found in the larval microbiomes were *Flavobacteriaceae*, *Saccharospirillaceae*, *Alteromonadaceae*, *Rhodobacteraceae*, and *Vibrionaceae* (Figure 2B, D-G). Among the microbiomes of larvae grown in LMR conditions, *Flavobacteriaceae* was present at all time points, but increased at 4 and 6 dpf (overall mean relative abundance 8% and 13%, respectively; Figure 2B). Similarly, when cultured in LMR, larval microbiomes exhibited increasing relative abundances of *Shewanellaceae* through development (overall mean relative abundance from 5% to 29%; Figure 2B), whereas *Saccharospirillaceae* was initially predominant in the larval microbiome but declined sharply from 2 to 6 dpf (overall mean relative abundance from 32% to 4%; Figure 2B, F). *Alteromonadaceae* displayed variable trends through development; abundance was greatest at 4 dpf, just after the first feeding, and declined at 6 dpf in all treatments except 18 °C LMR (20% overall mean relative abundance at 4 dpf; Figure 2B, D). In larvae at 18 °C LMR, *Alteromonadaceae* was most highly abundant at 6 dpf (29% overall mean relative abundance; Figure 2B, D). In 18 °C LMR, *Rhodobacteraceae* decreased in relative abundance throughout development (overall mean relative abundance from 24% to 7%; Figure 2B, E). Larvae cultured in 14 °C LMR had the greatest abundance of *Rhodobacteraceae* at 4 dpf (25% overall mean relative abundance; Figure 2B, E). In LMR conditions, *Vibrionaceae* generally increased from 2 to 4 dpf before stabilizing or decreasing by 6 dpf (14% overall mean relative abundance at 6 dpf; Figure 2B, G).

The microbiomes of larvae cultured in HMR conditions also maintained relative stability of families *Alteromonadaceae*, *Flavobacteriaceae*, and *Vibrionaceae* across developmental stages (Figure 2B, D, G). *Saccharospirillaceae* was nearly absent in the HMR larval microbiomes (3% overall mean relative abundance) (Figure 2B, F). In HMR conditions, regardless of temperature, *Rhodobacteraceae* exhibited the highest abundance at 2 dpf (17% overall mean relative abundance), sharply declined at 4 dpf (4%), and then slightly increased at 6 dpf (10%) (Figure 2B, E). Two families – *Crocinitomicaceae* and *Psychromonadaceae –* appeared in the HMR condition of larval microbiomes, but with low presence (5% and 3% overall mean relative abundance, respectively; Figure 2B). Overall, these family-specific successional patterns reveal that larval microbial communities shift throughout the pluteus stage.

### Larval microbiomes exhibit host and environmental selectivity

To determine if the larval microbiome is primarily shaped by environmental exposure or selective processes within the host, we compared the microbial communities within the larvae and the surrounding seawater at 6 dpf (Figure 3). Although the larval microbiomes contained microbial groups present in the seawater surrounding them, the relative abundances of these groups differed between the larvae and seawater. *Alteromonadaceae*, *Rhodobacteraceae*, and *Flavobacteriaceae* were found in both seawater and larvae; however, *Rhodobacteraceae* and *Flavobacteriaceae* were more prevalent in seawater, while *Alteromonadaceae* was more abundant in larvae (Figure 3A). *Saccharospirillaceae* and *Pseudoalteromonadaceae* were less abundant in both seawater and larvae than other taxa, regardless of temperature (Figure 3A). By 6 dpf, larval microbiomes in LMR conditions contained high relative abundances of *Vibrionaceae* (31% overall mean relative abundance) and *Shewanellaceae* (28%) at 14 °C (Figure 3A). Similarly, in 18 °C LMR, *Alteromonadaceae* (29%) and *Shewanellaceae* (30%) were abundant (Figure 3A). Regardless of temperature, both HMR seawater and larvae harbored high relative abundances of *Rhodobacteraceae* and *Flavobacteriaceae* (Figure 3A). Notably, a large portion of the larval microbiome in HMR conditions was composed of less abundant families categorized as “Other” (*i.e.*, families that comprised < 25% relative abundance in any single sample) (Figure 3A).

**Figure 3.**
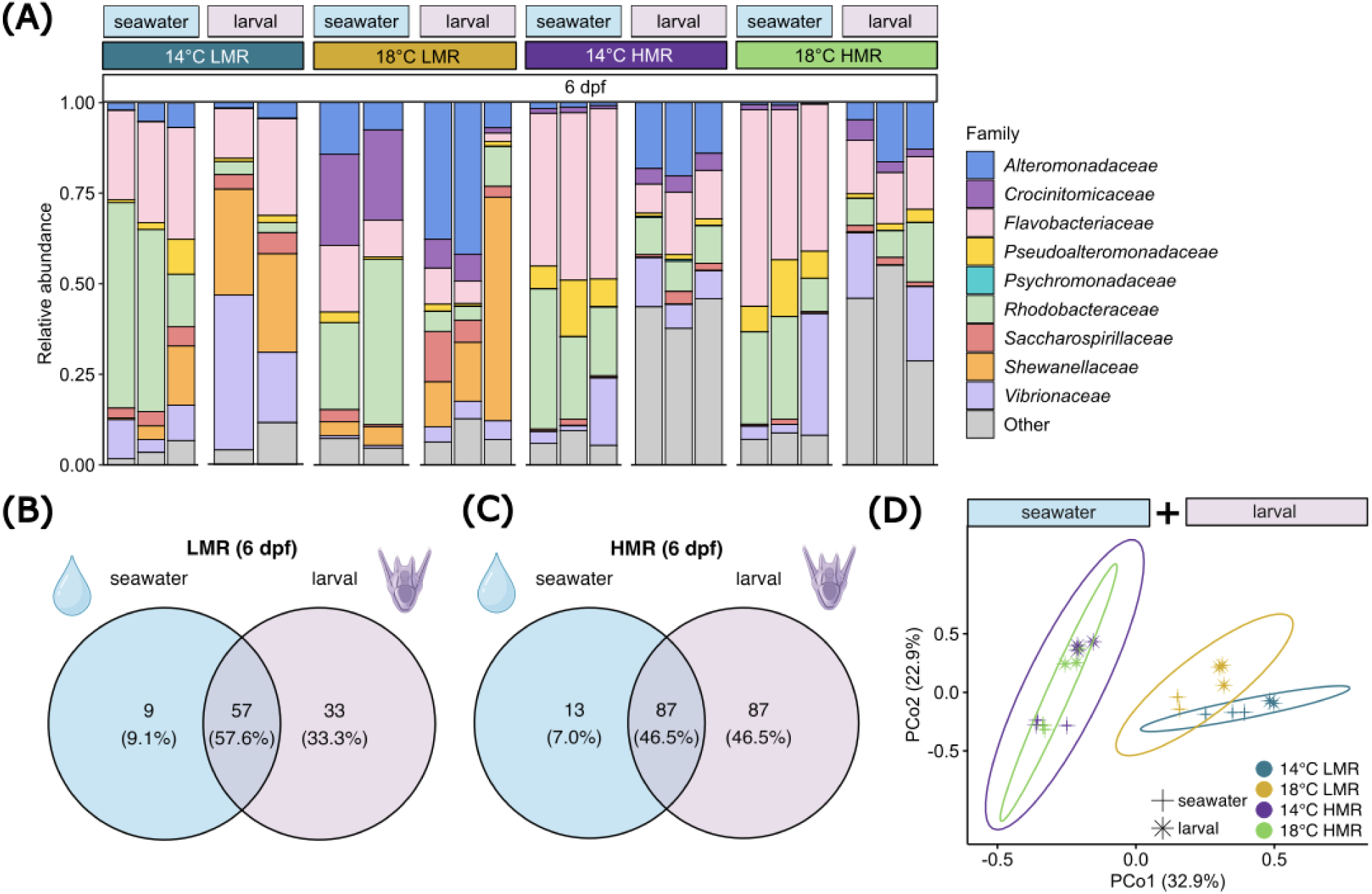
Larvae select their microbiome from the surrounding seawater. A) Stacked bar plot with relative abundance of seawater and larval microbial communities across four treatments at 6 days post-fertilization (dpf). Each bar represents a sample, with colors indicating the top nine families (≥ 25% relative abundance in any single sample). “Other” consists of the remaining families grouped as one category. (B and C) Venn diagrams of the number of families in low microbial richness (LMR) (B) and high microbial richness (HMR) (C) conditions in seawater and larval samples at 6 dpf. D).Bray-Curtis beta diversity across four treatments in seawater and larval samples at 6 dpf. Each point is a sample and shape indicates sample type (i.e., seawater and larvae).

LMR conditions promoted similar microbial communities in seawater and larvae, whereas HMR conditions led to more distinct larval microbiomes (Figure 3A-C). At 6 dpf, LMR conditions shared 57 (57.6%) families between seawater and larval samples, with 9 (9.1%) unique to seawater and 33 (33.3%) families unique to larvae (Figure 3B). Conversely, HMR conditions exhibited a much greater overlap between larvae and seawater, with 87 (46.5%) shared families and 87 (46.5%) families exclusive to larval microbiomes (Figure 3C). Beta diversity revealed that microbial richness (R^2^ = 0.31, *p*_PERMANOVA_ = 0.001) and the interaction of temperature and microbial richness (R^2^ = 0.07, *p*_PERMANOVA_ = 0.037) were significant, but temperature was not (R^2^ = 0.05, *p*_PERMANOVA_ = 0.147) (Figure 3D), with both LMR and HMR conditions exhibiting distinct clustering regardless of temperature, while seawater and larval samples also grouped separately (Figure 3D).

*ANCOM-BC* (Lin & Peddada, 2020) was used to analyze the differential abundance of taxa, revealing specific microbial taxa with the greatest changes based on temperature (18 °C vs. 14 °C) and microbial seawater condition (HMR vs. LMR). At 14 °C, genera *Celeribacter* (family *Rhodobacteraceae*) and *Alkalimarinus* (family *Alteromonadaceae*) were enriched relative to 18 °C in seawater samples (Figure 4A). *Neisseria* (family *Neisseriaceae*) was highly enriched in both seawater and larval microbial communities at 14 °C, relative to 18 °C (Figure 4A, B). Meanwhile, both seawater and larval microbiomes exhibited shared enrichment of genus *Nautella* (family *Rhodobacteraceae*) at 18 °C (Figure 4A, B). Multiple genera were significantly enriched in HMR (compared to LMR), such as *Lentibacter* (family *Rhodobacteraceae*) and *Aliivibrio* (family *Vibrionaceae*) in seawater (Figure 4C), and *Flavicella* (family *Flavobacteriaceae*) in larvae (Figure 4D). Contrarily, in LMR seawater and larvae, there was enrichment of *Ekhidna* (family *Reichenbachiellaceae*), *Psychrobium* (family *Shewanellaceae*), and *Bermanella* (family *Oceanospirillaceae*) (Figure 4C, D).

**Figure 4.**
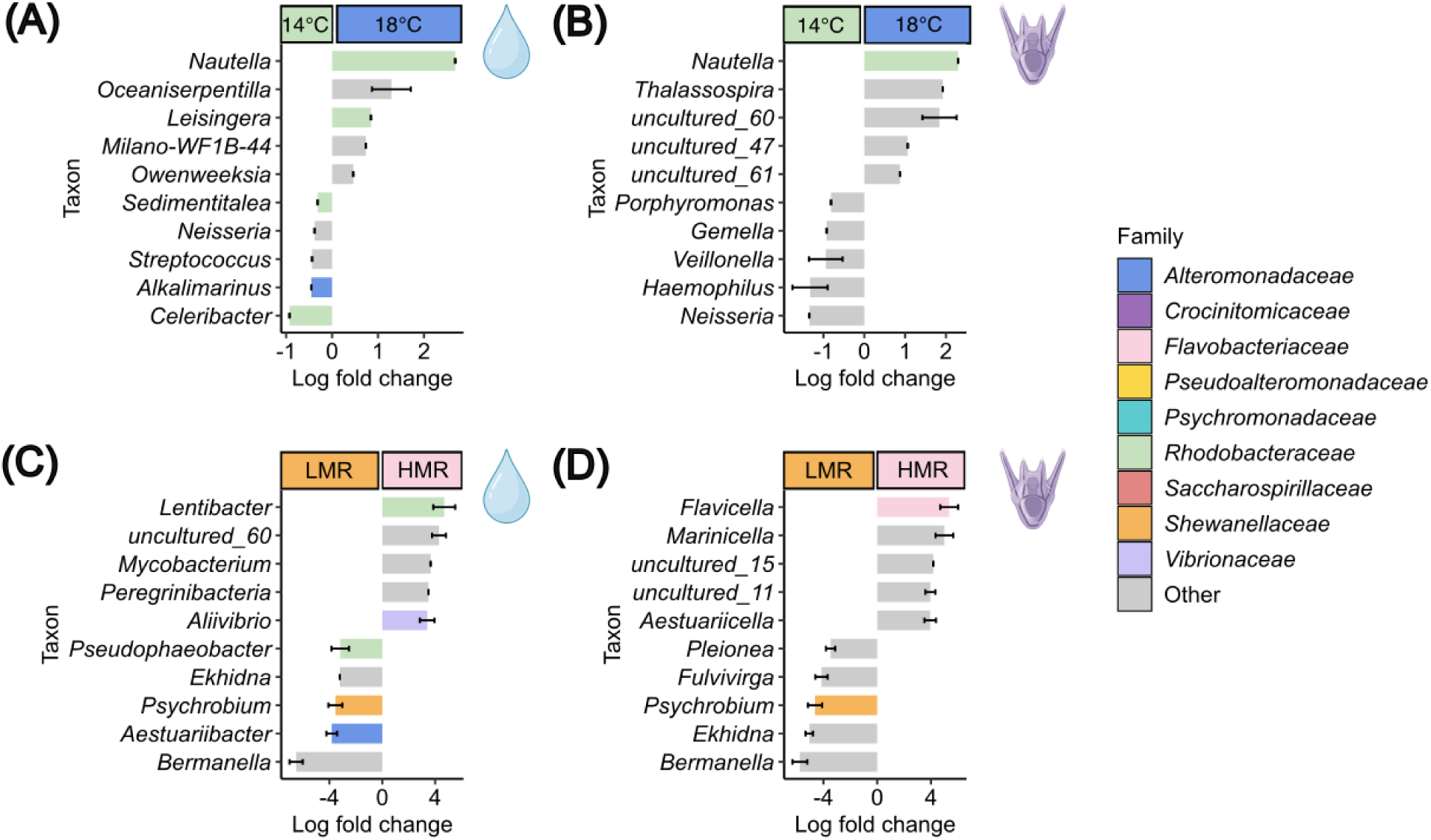
The most highly enriched microbes differ between larvae and their seawater environment. (A-B) Log fold changes of taxa enriched in seawater (A) and larvae (B) at all time points relative to 14 °C conditions. (C-D) Log fold changes of taxa enriched in seawater (C) and larvae (D) at all time points relative to low microbial richness (LMR) conditions. ‘Candidatus_Peregrinibacteria’ was shortened to ‘Peregrinibacteria’ for visualization. Each bar represents a genus, with colors indicating the family each genus belongs to out of the top nine families (≥ 25% relative abundance in any single sample). “Other” consists of the remaining families grouped as one category.

### Microbial-mediated influences on larval morphology

Larval microbiomes at 6 dpf were compared to the morphological results to determine if there was an association between host-associated microbiomes and larval growth (Figure 5A). Larvae grown in different microbial richness exhibit differences in morphology. At 6 dpf, larval length was significantly impacted by temperature (*p_ANOVA_* = 0.003), microbial richness (*p_ANOVA_* = 0.040), and the interaction of these two (temperature × seawater condition; *p_ANOVA_* = 0.024) (Figure 5B, C).

**Figure 5.**
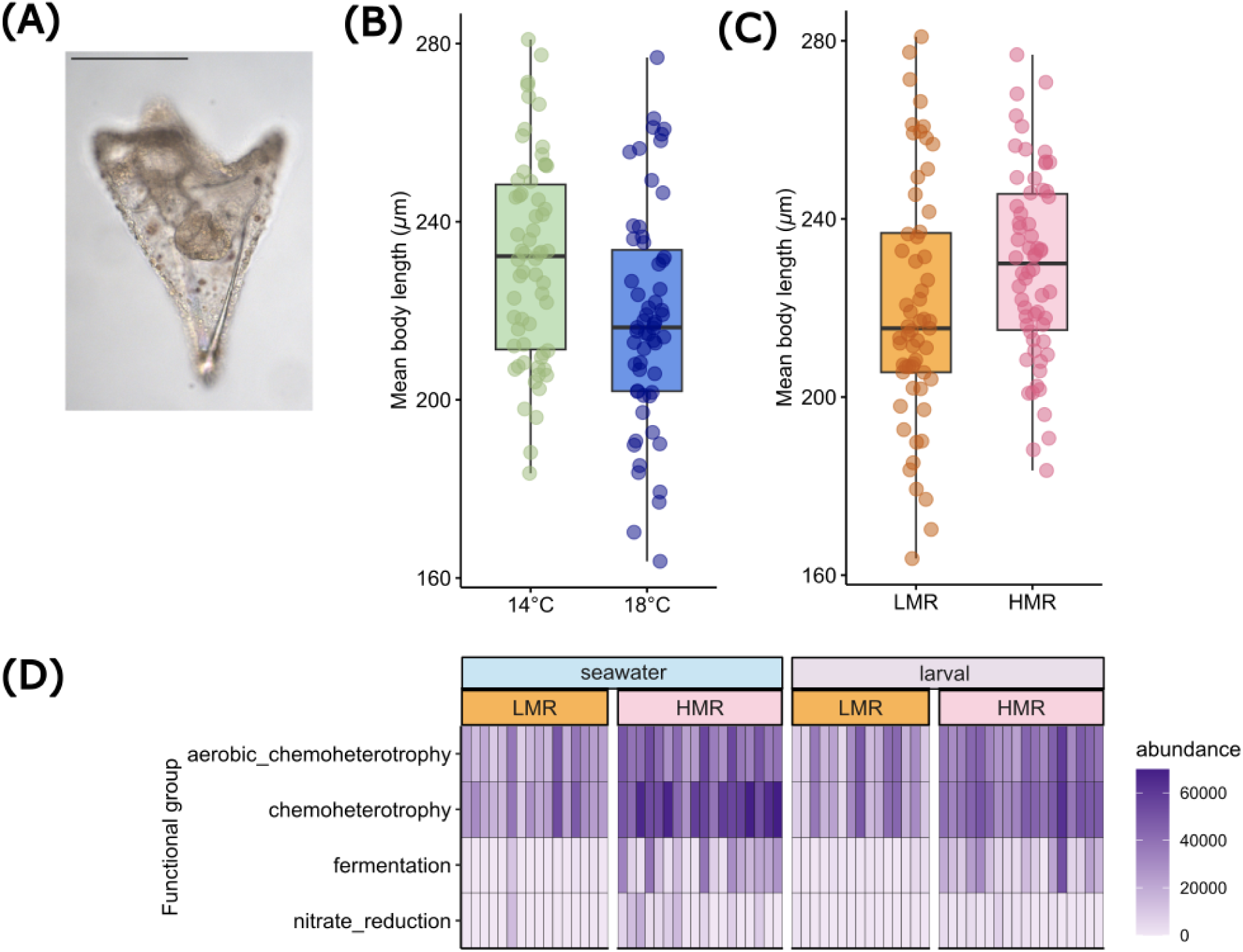
Developmental temperature, microbial communities, and their functions impact larval morphology. A) Representative larva at 6 days post-fertilization (dpf). Scale bar = 100 μm B) Larval mean body length across temperature at 6 dpf. Each point is a sample and each box represents the interquartile range (IQR) with the median line, whiskers extend to 1.5× the IQR. C) Larval mean body length across microbial seawater conditions at 6 dpf. Each point is a sample and each box represents the IQR with the median line, whiskers extend to 1.5× the IQR. D) Heatmaps of the predicted functional groups of microbes present in LMR and HMR conditions in seawater and larval samples at all time points. Each bar is a sample.

To investigate how the microbiomes of larvae and seawater could potentially metabolically influence larval growth, FAPROTAX (Louca et al., 2016) was used to predict the metabolic functions of the microbial communities in seawater and larval samples. Microbial communities in seawater and larvae had a high abundance of functions associated with ‘aerobic chemoheterotrophy’ and ‘chemoheterotrophy’ (Figure 5D), suggesting that the microbes primarily obtain their energy and carbon from organic compounds. Additionally, in HMR, regardless of temperature, there were predicted functions related to ‘fermentation’ and ‘nitrate reduction’ (Figure 5D).

## Discussion

### Temperature does not influence sea urchin larval microbiome diversity

Environmental conditions dictate not only which microbes are available to seed host microbiomes but can also directly influence established host-associated microbiomes (Griffiths et al., 2019; Pantos et al., 2015; Yang et al., 2024). Here, we investigated how seawater temperature and microbial richness shape the sea urchin larval microbiome throughout early development. We found that larval microbiome alpha and beta diversity were better explained by the microbial load and richness of the seawater, rather than temperature. This contrasts findings in other marine invertebrates where elevated temperature shifts microbiome composition, opening niches for opportunistic pathogens under heat stress (Li et al., 2022; Li et al., 2018; De Castro-Fernández et al., 2023; Ramsby et al., 2018; Ritchie, 2006). The lack of observed temperature effect on larval-associated microbiome diversity in our study may, in part, be due to the wide thermal tolerance of many marine bacterial taxa (Garcia et al., 2022; Rajeev et al., 2021). Since the HMR seawater was sourced from an aquarium tank maintained at 15 °C, the higher temperature of 18 °C might not have been a substantial enough temperature change to disrupt the established microbial communities. It is also possible that other environmental factors (*e.g.,* nutrient availability) could mask the effect of the elevated temperature (Wang et al., 2016). Alternatively, a robust host immune system could contribute to maintaining and stabilizing the larval microbiome at 18 °C (Nyholm & Graf, 2012; Taylor & Vega, 2021). In contrast, the heat stress disruption of microbiomes in other systems could be a consequence of a weakened immune system. Although temperature changes can alter environmental microbes and host microbiomes, the results presented here highlight the strong influence of microbial communities on the formation of the host-associated microbiome.

### Larval microbial communities exhibit some degree of host selectivity

Established microbiomes impact host development and physiology, including the development of the immune system (Fallet et al., 2022). The predominant microbial families identified in our seawater and larval samples – *Rhodobacteraceae*, *Flavobacteriaceae*, and *Alteromonadaceae* – are broadly distributed throughout the ocean and commonly found in marine invertebrate microbiomes, including early developmental stages of echinoderms (Haditomo et al., 2021; Marangon et al., 2023; Miller et al., 2021; Yu et al., 2023). Family *Rhodobacteraceae* are abundant in biofilms, making up to 30% of pelagic bacterial communities and contributing to biogeochemical cycling (Elifantz et al., 2013; Simon et al., 2017). *Flavobacteriaceae* play key roles in degrading polysaccharides (Mann et al., 2013; Zhou et al., 2016), whereas *Alteromonadaceae* are widespread chemoorganotrophs (Ivanova & Mikhailov, 2001). *Vibrionaceae* were also observed across all four treatments and, while likely involved in nutrient cycling (Dryselius et al., 2007), may represent opportunistic associations rather than stable or host-specific relationships (Ritchie, 2006; Takemura et al., 2014; Thompson et al., 2004). Overall, we find that the microbial communities present in our larval cultures are ubiquitous marine organisms known to form associations with marine invertebrates in the natural environment.

Selection shapes the assembly of the host microbiome in early development (Flores et al., 2025) and typically requires cell surface contact and cellular signaling between host and microbes, although exact methods of selection vary widely from mechanical to chemical signaling (Dunn & Weis, 2009; Nyholm & McFall-Ngai, 2021). We observed differences between the microbial content of the seawater and larvae, which supports previous observations that larval sea urchins select microbial communities (Carrier & Reitzel, 2018). Here, the different microbial profiles are particularly evident in the higher relative abundance of *Rhodobacteraceae* and *Flavobacteriaceae* within seawater and the higher abundance of *Alteromonadaceae* in larvae, supporting the idea of larval selectivity.

Larvae grown in HMR seawater hosted more microbial families than larvae in LMR, which is consistent with HMR seawater itself having a higher microbial load and diversity. Seawater and larvae shared a high percentage of microbial families at 2 dpf: 40.4% in LMR and 53.5% in HMR, showing that the environment provides the initial microbial communities available to the larvae. After initial uptake, however, the larval-associated microbiome undergoes selection or enrichment by the larval host, as evidenced by the higher alpha diversity in larvae under HMR conditions and the presence of taxa unique to larval samples. By 6 dpf, microbial families in seawater and larvae overlapped by 57.6% in LMR and 46.6% in HMR.

### Larval microbiomes shift across development

Winnowing refers to a selection process during early-life stages in which an initially highly diverse microbiome becomes fine-tuned during the course of development (Epstein et al., 2019; Nyholm & McFall-Ngai, 2004; Littman et al., 2009a; Littman et al., 2009b). Here, we did not observe microbiome winnowing across developmental stages in *S. purpuratus* larvae. Although alpha diversity decreased from 2 to 4 dpf in most treatments, it slightly increased again at 6 dpf. It is possible, however, that winnowing occurs in larval purple sea urchins at later developmental stages. Carrier and Reitzel (2019) reported shifts in microbial diversity and a decrease in the amount of *Psychromonas* from early development to post-metamorphic juvenile stages in *S. purpuratus* and two other echinoid species, although the changes were modest even across major developmental stages. Alternatively, it is possible that *S. purpuratus* larvae lack microbiome specificity when reared in artificial seawater used in laboratory conditions (Schuh et al., 2020). However, the differences in the relative abundance of microbial families between the seawater and larval microbiomes suggest that some selection occurred.

Host genotype can also strongly influence microbiome composition (Ahern et al., 2021; Benga et al., 2024; Qin et al., 2022), however, studies in adult sea urchins suggest that genotype appears to have limited, or at least far less, influence on the gut microbiome than other factors (Redelinghuys et al., 2025; Rodríguez-Barreras et al., 2024). In sea urchin larvae, the environment is mainly responsible for microbial community differences, rather than genotype (Carrier et al., 2019). Similarly, we found trends were consistent across the familial crosses in each treatment, suggesting that environmental conditions had a strong effect on the microbiome despite underlying genetic variation. While the environment appears to drive initial microbiome establishment, it is possible that host genotype could influence selection seen at later developmental stages.

While the larval microbiomes remained diverse throughout development, we did observe small microbial community shifts in the larvae cultured in HMR conditions, whereas the larval microbiomes in LMR conditions remained relatively stable over time. This may suggest that the microbial community was stable in the absence of environmental microbes that could disrupt or restructure the larval microbiome. The increased availability of microbes in HMR conditions could create more opportunities for shifts in the microbial community over time, resulting in the changes in larval microbiomes observed from 2 to 6 dpf. One potential source of changes in the microbiome is feeding (Carrier et al., 2021), which occurred on 3 and 5 dpf. However, in our laboratory, algae were washed in sterilized seawater to prevent bacterial growth, which minimizes this possibility. Microbial diversity could be important to larval survival, as diverse microbiomes often support an organism’s capacity to withstand environmental stress (Dickey et al., 2025; Pinnow et al., 2023).

### Larval morphology may be influenced by the developing microbiome

Our results show that larval size at 6 dpf was impacted by both temperature and microbial communities, such that larvae were longer when cultured at 14 °C or HMR conditions. This is consistent with previous observations demonstrating that temperature is a strong driver of larval growth in echinoderms (Rahman et al., 2007; Wilkins et al., 2024) and other invertebrates (Guiraud et al., 2021; Mesler & Mabry, 2024). However, other studies have shown that these temperature-induced differences may be stage-specific and no longer evident by the pluteus stage (Wong et al., 2019). The observation that larval microbiomes also shape morphology further emphasizes the importance of understanding these early associations (Tan et al., 2025).

Microbial metabolism may also influence larval phenotype by the secretion or uptake of specific metabolites. The microbial communities present in both seawater and larvae in both LMR and HMR conditions were enriched in species with putative metabolic functions related to ‘chemoheterotrophy,’ a broad trophic function that makes up a major role in marine and aquatic ecosystems (Cui et al., 2019; Guan et al., 2025; Amblard et al., 1992). Chemoheterotrophy contributes to active organic matter turnover and the carbon cycle (Passos et al., 2022), all of which can modify the chemical microenvironment surrounding larvae and the larvae themselves, thus having the potential to influence their morphology.

In contrast, the microbial communities characterized in the HMR seawater and larval microbiomes were enriched in species associated with ‘fermentation,’ which likely reflects a greater metabolic activity. This may increase oxygen demand and accelerate nutrient cycling, creating a more dynamic environment. Because bacterial fermentation produces metabolites that include short-chain fatty acids, enrichment of fermentation pathways suggests production of these metabolites (Rowland et al., 2018). Additionally, ‘nitrate reduction’ was enriched in the strains identified in HMR seawater, potentially explaining some of the observed larval size differences. Furthermore, previous work suggests that a stable gut microbiome enhances nutrient metabolic efficiency (Kong et al., 2024), which might be reflected in other observed variations in larval morphology patterns. Alternatively, trade-offs may exist between microbiome selection and growth. Since selection is an active process by the host, it requires expenditure of resources via the production of metabolites, antimicrobial compounds, or immune system regulation (Flores et al., 2025). Functions such as ‘chemoheterotrophy,’ ‘fermentation,’ and ‘nitrate reduction’ were also found in the microbiomes of juvenile Antarctic whelks (*Neobuccinum eatoni*), and likely aid in digestion (Buschi et al., 2025). Further, Haditomo et al. (2021) suggested that nitrate in the sea urchin gut microbiome is reduced to ammonium, potentially enhancing host growth. Here, we found ‘nitrate reduction’ to be primarily enriched in seawater samples but not larval microbiomes, which still have the ability to indirectly impact larvae by altering the chemical environment surrounding them. Consequently, this interaction between the developing larval microbiome and growth has the potential to impact their survival, as larval size impacts resource acquisition and predation risk (Allen, 2008; Dingeldein & White, 2016; Garrido et al., 2015). Together, these findings suggest that the microbes present in both seawater and larval samples could indirectly provide nutrients to the larvae through their byproducts, impacting larval morphology.

## Acknowledgements

We would like to formally acknowledge the work that contributed to this publication: Emily Wilkins, Will Kotas, and Grace Mathis for imaging larvae for morphometric analyses; Mikayla Clark and Grace Mathis for assessing bacterial load using qPCR; and Sidney Pasternak, Sarah Gonzalez Martinez, and Megan Anglin for generating larval morphometric measurements. Computational analysis was conducted with the advanced computing resources provided by Texas A&M High Performance Research Computing.

## Data Availability Statement

Raw sequence reads and associated metadata are deposited in the SRA (BioProject PRJNA1447470). Scripts and code used for analysis can be found in GitHub repository: https://github.com/stephaniehendricks/MM16S.

## References

1. Abdelfadil, M. R., Patz, S., Kolb, S., & Ruppel, S. (2024). Unveiling the influence of salinity on bacterial microbiome assembly of halophytes and crops. Environmental Microbiome, 19(1), 49. 10.1186/s40793-024-00592-3

2. Ahern, O. M., Whittaker, K. A., Williams, T. C., Hunt, D. E., & Rynearson, T. A. (2021). Host genotype structures the microbiome of a globally dispersed marine phytoplankton. Proceedings of the National Academy of Sciences, 118(48), e2105207118. 10.1073/pnas.2105207118

3. Allen, J. D. (2008). Size-Specific Predation on Marine Invertebrate Larvae. The Biological Bulletin, 214(1), 42–49. 10.2307/25066658

4. Anderson, M. J. (2001). A new method for non-parametric multivariate analysis of variance.

5. Angthong, P., Uengwetwanit, T., Arayamethakorn, S., Chaitongsakul, P., Karoonuthaisiri, N., & Rungrassamee, W. (2020). Bacterial analysis in the early developmental stages of the black tiger shrimp (Penaeus monodon). Scientific Reports, 10(1), 4896. 10.1038/s41598-020-61559-1

6. Archie, E. A., & Tung, J. (2015). Social behavior and the microbiome. Current Opinion in Behavioral Sciences, 6, 28–34. 10.1016/j.cobeha.2015.07.008

7. Baquiran, J. I. P., Quijano, J. B., Van Oppen, M. J. H., Cabaitan, P. C., Harrison, P. L., & Conaco, C. (2025). Microbiome Stability Is Linked to *Acropora* Coral Thermotolerance in Northwestern Philippines. Environmental Microbiology, 27(2), e70041. 10.1111/1462-2920.70041

8. Battaglia, T. (2018). btools: A suite of R function for all types of microbial diversity analyses. (Version 0.0.1) [Computer software]. https://rdrr.io/github/twbattaglia/btools/

9. Benga, L., Rehm, A., Gougoula, C., Westhoff, P., Wachtmeister, T., Benten, W. P. M., Engelhardt, E., Weber, A. P. M., Köhrer, K., Sager, M., & Janssen, S. (2024). The host genotype actively shapes its microbiome across generations in laboratory mice. Microbiome, 12(1), 256. 10.1186/s40168-024-01954-2

10. Bengtsson, M. M., Helgesen, M., Wang, H., Fredriksen, S., & Norderhaug, K. M. (2025). Sea urchin intestinal bacterial communities depend on seaweed diet and contain nitrogen-fixing symbionts. FEMS Microbiology Ecology, 101(2), fiaf006. 10.1093/femsec/fiaf006

11. Bisanz, J. (2018). qiime2R: Importing QIIME2 artifacts and associated data into R sessions (Version v0.99) [Computer software]. https://github.com/jbisanz/qiime2R

12. Bolyen, E., Rideout, J. R., Dillon, M. R., Bokulich, N. A., Abnet, C. C., Al-Ghalith, G. A., Alexander, H., Alm, E. J., Arumugam, M., Asnicar, F., Bai, Y., Bisanz, J. E., Bittinger, K., Brejnrod, A., Brislawn, C. J., Brown, C. T., Callahan, B. J., Caraballo-Rodríguez, A. M., Chase, J., … Caporaso, J. G. (2019). Reproducible, interactive, scalable and extensible microbiome data science using QIIME 2. Nature Biotechnology, 37(8), 852–857. 10.1038/s41587-019-0209-9

13. Bordenstein, S. R., & Theis, K. R. (2015). Host Biology in Light of the Microbiome: Ten Principles of Holobionts and Hologenomes. PLOS Biology, 13(8), e1002226. 10.1371/journal.pbio.1002226

14. Bright, M., & Bulgheresi, S. (2010). A complex journey: Transmission of microbial symbionts. Nature Reviews Microbiology, 8(3), 218–230. 10.1038/nrmicro2262

15. Brodeur, R. D., Auth, T. D., & Phillips, A. J. (2019). Major Shifts in Pelagic Micronekton and Macrozooplankton Community Structure in an Upwelling Ecosystem Related to an Unprecedented Marine Heatwave. Frontiers in Marine Science, 6, 212. 10.3389/fmars.2019.00212

16. Burton, T., & Metcalfe, N. B. (2014). Can environmental conditions experienced in early life influence future generations? Proceedings of the Royal Society B: Biological Sciences, 281(1785), 20140311. 10.1098/rspb.2014.0311

17. Buschi, E., Tangherlini, M., Martire, M. L., & Corinaldesi, C. (2025). The juvenile Antarctic whelk Neobuccinum eatoni maintains a specialized microbiome in its proboscis even in adulthood. Polar Biology, 48(2), 68. 10.1007/s00300-025-03388-4

18. Carrier, T. J., Dupont, S., & Reitzel, A. M. (2019). Geographic location and food availability offer differing levels of influence on the bacterial communities associated with larval sea urchins. FEMS Microbiology Ecology, 95(8), fiz103. 10.1093/femsec/fiz103

19. Carrier, T. J., & Reitzel, A. M. (2018). Convergent shifts in host-associated microbial communities across environmentally elicited phenotypes. Nature Communications, 9(1), 952. 10.1038/s41467-018-03383-w

20. Carrier, T., & Reitzel, A. (2019). Bacterial community dynamics during embryonic and larval development of three confamilial echinoids. Marine Ecology Progress Series, 611, 179–188. 10.3354/meps12872

21. Cavalcanti, G. S., Alker, A. T., Delherbe, N., Malter, K. E., & Shikuma, N. J. (2020). The Influence of Bacteria on Animal Metamorphosis. Annual Review of Microbiology, 74(1), 137–158. 10.1146/annurev-micro-011320-012753

22. Chen, X., Mo, L., Zhang, L., Huang, L., Gao, Z., Peng, J., Yu, Z., & Zhang, X. (2024). Taxonomic Diversity, Predicted Metabolic Pathway, and Interaction Pattern of Bacterial Community in Sea Urchin Anthocidaris crassispina. Microorganisms, 12(10), 2094. 10.3390/microorganisms12102094

23. Chuang, P.-S., Yu, S.-P., Liu, P.-Y., Hsu, M.-T., Chiou, Y.-J., Lu, C.-Y., & Tang, S.-L. (2024). A gauge of coral physiology: Re-examining temporal changes in *Endozoicomonas* abundance correlated with natural coral bleaching. ISME Communications, 4(1), ycae001. 10.1093/ismeco/ycae001

24. Comizzoli, P., Power, M. L., Bornbusch, S. L., & Muletz-Wolz, C. R. (2021). Interactions between reproductive biology and microbiomes in wild animal species. Animal Microbiome, 3(1), 87. 10.1186/s42523-021-00156-7

25. De Castro-Fernández, P., Ballesté, E., Angulo-Preckler, C., Biggs, J., Avila, C., & García-Aljaro, C. (2023). How does heat stress affect sponge microbiomes? Structure and resilience of microbial communities of marine sponges from different habitats. Frontiers in Marine Science, 9, 1072696. 10.3389/fmars.2022.1072696

26. Dickey, J. R., Mercer, N. M., Kuijpers, M. C. M., Props, R., & Jackrel, S. L. (2025). Biodiversity within phytoplankton-associated microbiomes regulates host physiology, host community ecology, and nutrient cycling. mSystems, 10(2), e01462–24. 10.1128/msystems.01462-24

27. Dingeldein, A. L., & White, J. W. (2016). Larval traits carry over to affect post-settlement behaviour in a common coral reef fish. Journal of Animal Ecology, 85(4), 903–914. 10.1111/1365-2656.12506

28. Donia, M. S., Fricke, W. F., Partensky, F., Cox, J., Elshahawi, S. I., White, J. R., Phillippy, A. M., Schatz, M. C., Piel, J., Haygood, M. G., Ravel, J., & Schmidt, E. W. (2011). Complex microbiome underlying secondary and primary metabolism in the tunicate-*Prochloron* symbiosis. Proceedings of the National Academy of Sciences, 108(51). 10.1073/pnas.1111712108

29. Eglaine, Z., Courboulès, J., Cipolletta, F., Roques, C., Soulié, T., Sarno, D., Mostajir, B., & Vidussi, F. (2025). Effects of a simulated marine heatwave on the structure and composition of Mediterranean plankton in a mesocosm study. PLOS One, 20(11), e0337112. 10.1371/journal.pone.0337112

30. Elkin, L. A., Kay, M., Higgins, J. J., & Wobbrock, J. O. (2021). An Aligned Rank Transform Procedure for Multifactor Contrast Tests. The 34th Annual ACM Symposium on User Interface Software and Technology, 754–768. 10.1145/3472749.3474784

31. Engel, P., & Moran, N. A. (2013). The gut microbiota of insects – diversity in structure and function. FEMS Microbiology Reviews, 37(5), 699–735. 10.1111/1574-6976.12025

32. Epstein, H. E., Torda, G., Munday, P. L., & Van Oppen, M. J. H. (2019). Parental and early life stage environments drive establishment of bacterial and dinoflagellate communities in a common coral. The ISME Journal, 13(6), 1635–1638. 10.1038/s41396-019-0358-3

33. Fallet, M., Montagnani, C., Petton, B., Dantan, L., De Lorgeril, J., Comarmond, S., Chaparro, C., Toulza, E., Boitard, S., Escoubas, J.-M., Vergnes, A., Le Grand, J., Bulla, I., Gueguen, Y., Vidal-Dupiol, J., Grunau, C., Mitta, G., & Cosseau, C. (2022). Early life microbial exposures shape the Crassostrea gigas immune system for lifelong and intergenerational disease protection. Microbiome, 10(1), 85. 10.1186/s40168-022-01280-5

34. Flores, C., Millard, S., & Seekatz, A. M. (2025). Bridging Ecology and Microbiomes: Applying Ecological Theories in Host-associated Microbial Ecosystems. Current Clinical Microbiology Reports, 12(1), 9. 10.1007/s40588-025-00246-z

35. Fox, J., & Weisberg, S. (2019). *An R Companion to Applied Regression* (Version 3.1-5) [Computer software]. 10.32614/CRAN.package.car

36. French, K. B., Herrera, M. J., & German, D. P. (2024). Sea Urchin Larvae (*Strongylocentrotus purpuratus*) Select and Maintain a Unique Microbiome Compared to Environmental Sources. The Biological Bulletin, 247(1), 56–73. 10.1086/736931

37. Garcia, F. C., Warfield, R., & Yvon-Durocher, G. (2022). Thermal traits govern the response of microbial community dynamics and ecosystem functioning to warming. Frontiers in Microbiology, 13, 906252. 10.3389/fmicb.2022.906252

38. Garrido, S., Ben-Hamadou, R., Santos, A. M. P., Ferreira, S., Teodósio, M. A., Cotano, U., Irigoien, X., Peck, M. A., Saiz, E., & Ré, P. (2015). Born small, die young: Intrinsic, size-selective mortality in marine larval fish. Scientific Reports, 5(1), 17065. 10.1038/srep17065

39. Grieneisen, L., Dasari, M., Gould, T. J., Björk, J. R., Grenier, J.-C., Yotova, V., Jansen, D., Gottel, N., Gordon, J. B., Learn, N. H., Gesquiere, L. R., Wango, T. L., Mututua, R. S., Warutere, J. K., Siodi, L., Gilbert, J. A., Barreiro, L. B., Alberts, S. C., Tung, J., … Blekhman, R. (2021). Gut microbiome heritability is nearly universal but environmentally contingent. Science, 373(6551), 181–186. 10.1126/science.aba5483

40. Griffiths, S. M., Antwis, R. E., Lenzi, L., Lucaci, A., Behringer, D. C., Butler, M. J., & Preziosi, R. F. (2019). Host genetics and geography influence microbiome composition in the sponge *Ircinia campana*. Journal of Animal Ecology, 88(11), 1684–1695. 10.1111/1365-2656.13065

41. Guiraud, M., Cariou, B., Henrion, M., Baird, E., & Gérard, M. (2021). Higher developmental temperature increases queen production and decreases worker body size in the bumblebee Bombus terrestris. Journal of Hymenoptera Research, 88, 39–49. 10.3897/jhr.88.73532

42. Haditomo, A. H. C., Yonezawa, M., Yu, J., Mino, S., Sakai, Y., & Sawabe, T. (2021). The Structure and Function of Gut Microbiomes of Two Species of Sea Urchins, Mesocentrotus nudus and Strongylocentrotus intermedius, in Japan. Frontiers in Marine Science, 8, 802754. 10.3389/fmars.2021.802754

43. Hakim, J. A., Schram, J. B., Galloway, A. W. E., Morrow, C. D., Crowley, M. R., Watts, S. A., & Bej, A. K. (2019). The Purple Sea Urchin Strongylocentrotus purpuratus Demonstrates a Compartmentalization of Gut Bacterial Microbiota, Predictive Functional Attributes, and Taxonomic Co-Occurrence. Microorganisms, 7(2), 35. 10.3390/microorganisms7020035

44. Hammer, T. J., Janzen, D. H., Hallwachs, W., Jaffe, S. P., & Fierer, N. (2017). Caterpillars lack a resident gut microbiome. Proceedings of the National Academy of Sciences, 114(36), 9641–9646. 10.1073/pnas.1707186114

45. Hartig, F., Lohse, L., & de Souza leite, M. (2024). DHARMa: Residual Diagnostics for Hierarchical (Multi-Level / Mixed) Regression Models (Version 0.4.7) [Computer software]. 10.32614/CRAN.package.DHARMa

46. Henry, L. P., Bruijning, M., Forsberg, S. K. G., & Ayroles, J. F. (2021). The microbiome extends host evolutionary potential. Nature Communications, 12(1), 5141. 10.1038/s41467-021-25315-x

47. Kay, M., Elkin, L., Higgins, J., & Wobbrock, J. (2025). *ARTool: Aligned RankTransform for Nonparametric Factorial ANOVAs* (Version 0.11.2) [Computer software]. 10.5281/zenodo.594511

48. Koch, H., & Schmid-Hempel, P. (2011). Socially transmitted gut microbiota protect bumble bees against an intestinal parasite. Proceedings of the National Academy of Sciences, 108(48), 19288–19292. 10.1073/pnas.1110474108

49. Kong, M., Zhao, W., Wang, C., Qi, J., Liu, J., & Zhang, Q. (2024). A Well-Established Gut Microbiota Enhances the Efficiency of Nutrient Metabolism and Improves the Growth Performance of Trachinotus ovatus. International Journal of Molecular Sciences, 25(10), 5525. 10.3390/ijms25105525

50. Koskella, B., & Bergelson, J. (2020). The study of host–microbiome (co)evolution across levels of selection. Philosophical Transactions of the Royal Society B: Biological Sciences, 375(1808), 20190604. 10.1098/rstb.2019.0604

51. Kuznetsova, A., Brockhoff, P. B., & Christensen, R. H. B. (2017). **lmerTest** Package: Tests in Linear Mixed Effects Models. Journal of Statistical Software, 82(13). 10.18637/jss.v082.i13

52. Kwong, W. K., & Moran, N. A. (2016). Gut microbial communities of social bees. Nature Reviews Microbiology, 14(6), 374–384. 10.1038/nrmicro.2016.43

53. Li, S., Qian, Z., Gao, S., Shen, W., Li, X., Li, H., & Chen, L. (2022). Effect of long-term temperature stress on the intestinal microbiome of an invasive snail. Frontiers in Microbiology, 13, 961502. 10.3389/fmicb.2022.961502

54. Li, Y.-F., Yang, N., Liang, X., Yoshida, A., Osatomi, K., Power, D., Batista, F. M., & Yang, J.-L. (2018). Elevated Seawater Temperatures Decrease Microbial Diversity in the Gut of Mytilus coruscus. Frontiers in Physiology, 9, 839. 10.3389/fphys.2018.00839

55. Lima, L. F. O., Weissman, M., Reed, M., Papudeshi, B., Alker, A. T., Morris, M. M., Edwards, R. A., De Putron, S. J., Vaidya, N. K., & Dinsdale, E. A. (2020). Modeling of the Coral Microbiome: The Influence of Temperature and Microbial Network. mBio, 11(2), e02691–19. 10.1128/mBio.02691-19

56. Lin, H., & Peddada, S. D. (2020). Analysis of compositions of microbiomes with bias correction. Nature Communications, 11(1), 3514. 10.1038/s41467-020-17041-7

57. Louca, S., Parfrey, L. W., & Doebeli, M. (2016). Decoupling function and taxonomy in the global ocean microbiome. Science, 353(6305), 1272–1277. 10.1126/science.aaf4507

58. Marangon, E., Uthicke, S., Patel, F., Marzinelli, E. M., Bourne, D. G., Webster, N. S., & Laffy, P. W. (2023). Life-stage specificity and cross-generational climate effects on the microbiome of a tropical sea urchin (Echinodermata: Echinoidea). Molecular Ecology, 32(20), 5645–5660. 10.1111/mec.17124

59. Martinson, V. G., Danforth, B. N., Minckley, R. L., Rueppell, O., Tingek, S., & Moran, N. A. (2011). A simple and distinctive microbiota associated with honey bees and bumble bees: THE MICROBIOTA OF HONEY BEES AND BUMBLE BEES. Molecular Ecology, 20(3), 619–628. 10.1111/j.1365-294X.2010.04959.x

60. Mazel, F., Davis, K. M., Loudon, A., Kwong, W. K., Groussin, M., & Parfrey, L. W. (2018). Is Host Filtering the Main Driver of Phylosymbiosis across the Tree of Life? mSystems, 3(5), 10.1128/msystems.00097-18. 10.1128/msystems.00097-18

61. McArdle, B. H., & Anderson, M. J. (2001). FITTING MULTIVARIATE MODELS TO COMMUNITY DATA: A COMMENT ON DISTANCE-BASED REDUNDANCY ANALYSIS. Ecology, 82(1), 290–297. 10.1890/0012-9658(2001)082%5B0290:FMMTCD%5D2.0.CO;2

62. McFall-Ngai, M., Hadfield, M. G., Bosch, T. C. G., Carey, H. V., Domazet-Lošo, T., Douglas, A. E., Dubilier, N., Eberl, G., Fukami, T., Gilbert, S. F., Hentschel, U., King, N., Kjelleberg, S., Knoll, A. H., Kremer, N., Mazmanian, S. K., Metcalf, J. L., Nealson, K., Pierce, N. E., … Wernegreen, J. J. (2013). Animals in a bacterial world, a new imperative for the life sciences. Proceedings of the National Academy of Sciences, 110(9), 3229–3236. 10.1073/pnas.1218525110

63. McMurdie, P. J., & Holmes, S. (2013). phyloseq: An R Package for Reproducible Interactive Analysis and Graphics of Microbiome Census Data. PLoS ONE, 8(4), e61217. 10.1371/journal.pone.0061217

64. Mesler, S. P., & Mabry, K. E. (2024). Effects of temperature experienced across life stages on morphology and flight behavior of painted lady butterflies (Vanessa cardui). Movement Ecology, 12(1), 76. 10.1186/s40462-024-00516-3

65. Miller, P. M., Lamy, T., Page, H. M., & Miller, R. J. (2021). Sea urchin microbiomes vary with habitat and resource availability. Limnology and Oceanography Letters, 6(3), 119–126. 10.1002/lol2.10189

66. Mogen, S. C., Lovenduski, N. S., Dallmann, A. R., Gregor, L., Sutton, A. J., Bograd, S. J., Quiros, N. C., Di Lorenzo, E., Hazen, E. L., Jacox, M. G., Buil, M. P., & Yeager, S. (2022). Ocean Biogeochemical Signatures of the North Pacific Blob. Geophysical Research Letters, 49(9), e2021GL096938. 10.1029/2021GL096938

67. Muffett, K. M., Labonté, J. M., & Miglietta, M. P. (2025). Florida Keys Cassiopea host benthos-like external microbiomes and a gut dominated by Vibrio, Endozoicomonas and Mycoplasma. PLOS One, 20(8), e0330180. 10.1371/journal.pone.0330180

68. Murray, M. G., & Thompson, W. F. (1980). Camegie Institution of Washington, Department of Plant Biology, Stanford, CA 94305, USA. Nucleic Acids Research.

69. Nyholm, S. V., & Graf, J. (2012). Knowing your friends: Invertebrate innate immunity fosters beneficial bacterial symbioses. Nature Reviews Microbiology, 10(12), 815–827. 10.1038/nrmicro2894

70. Nyholm, S. V., & McFall-Ngai, M. (2004). The winnowing: Establishing the squid–vibrio symbiosis. Nature Reviews Microbiology, 2(8), 632–642. 10.1038/nrmicro957

71. Nyholm, S. V., & McFall-Ngai, M. J. (2021). A lasting symbiosis: How the Hawaiian bobtail squid finds and keeps its bioluminescent bacterial partner. Nature Reviews Microbiology, 19(10), 666–679. 10.1038/s41579-021-00567-y

72. Oliver, E. C. J., Donat, M. G., Burrows, M. T., Moore, P. J., Smale, D. A., Alexander, L. V., Benthuysen, J. A., Feng, M., Sen Gupta, A., Hobday, A. J., Holbrook, N. J., Perkins-Kirkpatrick, S. E., Scannell, H. A., Straub, S. C., & Wernberg, T. (2018). Longer and more frequent marine heatwaves over the past century. Nature Communications, 9(1), 1324. 10.1038/s41467-018-03732-9

73. Pantos, O., Bongaerts, P., Dennis, P. G., Tyson, G. W., & Hoegh-Guldberg, O. (2015). Habitat-specific environmental conditions primarily control the microbiomes of the coral *Seriatopora hystrix*. The ISME Journal, 9(9), 1916–1927. 10.1038/ismej.2015.3

74. Park, J.-Y., Jo, J.-W., An, Y.-J., Lee, J.-J., & Kim, B.-S. (2023). Alterations in sea urchin (Mesocentrotus nudus) microbiota and their potential contributions to host according to barren severity. Npj Biofilms and Microbiomes, 9(1), 83. 10.1038/s41522-023-00450-z

75. Passos, J. G., Soares, L. F., Sumida, P. Y. G., Bendia, A. G., Nakamura, F. M., Pellizari, V. H., & Signori, C. N. (2022). Contribution of chemoautotrophy and heterotrophy to the microbial carbon cycle in the Southwestern Atlantic Ocean. Ocean and Coastal Research, 70(suppl 2), e22044. 10.1590/2675-2824070.22137jgp

76. Pepke, M. L., Hansen, S. B., & Limborg, M. T. (2024). Unraveling host regulation of gut microbiota through the epigenome–microbiome axis. Trends in Microbiology, 32(12), 1229–1240. 10.1016/j.tim.2024.05.006

77. Pinnow, N., Chibani, C. M., Güllert, S., & Weiland-Bräuer, N. (2023). Microbial community changes correlate with impaired host fitness of Aurelia aurita after environmental challenge. Animal Microbiome, 5(1), 45. 10.1186/s42523-023-00266-4

78. Pruesse, E., Quast, C., Knittel, K., Fuchs, B. M., Ludwig, W., Peplies, J., & Glockner, F. O. (2007). SILVA: A comprehensive online resource for quality checked and aligned ribosomal RNA sequence data compatible with ARB. Nucleic Acids Research, 35(21), 7188–7196. 10.1093/nar/gkm864

79. Qin, Y., Havulinna, A. S., Liu, Y., Jousilahti, P., Ritchie, S. C., Tokolyi, A., Sanders, J. G., Valsta, L., Brożyńska, M., Zhu, Q., Tripathi, A., Vázquez-Baeza, Y., Loomba, R., Cheng, S., Jain, M., Niiranen, T., Lahti, L., Knight, R., Salomaa, V., … Méric, G. (2022). Combined effects of host genetics and diet on human gut microbiota and incident disease in a single population cohort. Nature Genetics, 54(2), 134–142. 10.1038/s41588-021-00991-z

80. Quast, C., Pruesse, E., Yilmaz, P., Gerken, J., Schweer, T., Yarza, P., Peplies, J., & Glöckner, F. O. (2012). The SILVA ribosomal RNA gene database project: Improved data processing and web-based tools. Nucleic Acids Research, 41(D1), D590–D596. 10.1093/nar/gks1219

81. Quigley, K. M., Alvarez-Roa, C., Raina, J.-B., Pernice, M., & Van Oppen, M. J. H. (2023). Heat-evolved microalgal symbionts increase thermal bleaching tolerance of coral juveniles without a trade-off against growth. Coral Reefs, 42(6), 1227–1232. 10.1007/s00338-023-02426-z

82. R Core Team. (2023). R: A Language and Environment for Statistical Computing (Version 4.3.1) [Computer software]. https://www.R-project.org/

83. Rahman Md. S., Rahman Sk. M., & Uehara T. (2007). Effects of temperature on early development of the sea urchin Echinometra mathaei from the intertidal reef of Okinawa Island, Japan. Journal of the Japanese Coral Reef Society, 9(1), 35–48. 10.3755/jcrs.9.35

84. Rajeev, M., Sushmitha, T. J., Aravindraja, C., Toleti, S. R., & Pandian, S. K. (2021). Thermal discharge-induced seawater warming alters richness, community composition and interactions of bacterioplankton assemblages in a coastal ecosystem. Scientific Reports, 11(1), 17341. 10.1038/s41598-021-96969-2

85. Ramsby, B. D., Hoogenboom, M. O., Whalan, S., & Webster, N. S. (2018). Elevated seawater temperature disrupts the microbiome of an ecologically important bioeroding sponge. Molecular Ecology, 27(8), 2124–2137. 10.1111/mec.14544

86. Redelinghuys, S., Emami-Khoyi, A., Matcher, G., Teske, P. R., Heltai, M., Csányi, S., Toonen, R. J., & Porri, F. (2025). Gut Microbial Diversity and Genome-Wide Variation of the Cape Sea Urchin (*Parechinus angulosus*) Across a Thermal Gradient. Austral Ecology, 50(10), e70118. 10.1111/aec.70118

87. Ritchie, K. (2006). Regulation of microbial populations by coral surface mucus and mucus-associated bacteria. Marine Ecology Progress Series, 322, 1–14. 10.3354/meps322001

88. Robeson, M. S., O’Rourke, D. R., Kaehler, B. D., Ziemski, M., Dillon, M. R., Foster, J. T., & Bokulich, N. A. (2021). RESCRIPt: Reproducible sequence taxonomy reference database management. PLOS Computational Biology, 17(11), e1009581. 10.1371/journal.pcbi.1009581

89. Rodríguez-Barreras, R., Tosado-Rodríguez, E. L., Dominicci-Maura, A., & Godoy-Vitorino, F. (2024). Effects of temperature and size class on the gut digesta microbiota of the sea urchin *Tripneustes ventricosus*. PeerJ, 12, e18298. 10.7717/peerj.18298

90. Rodríguez-Barreras, R., Tosado-Rodríguez, E. L., & Godoy-Vitorino, F. (2021). Trophic niches reflect compositional differences in microbiota among Caribbean sea urchins. PeerJ, 9, e12084. 10.7717/peerj.12084

91. Rowland, I., Gibson, G., Heinken, A., Scott, K., Swann, J., Thiele, I., & Tuohy, K. (2018). Gut microbiota functions: Metabolism of nutrients and other food components. European Journal of Nutrition, 57(1), 1–24. 10.1007/s00394-017-1445-8

92. Sadeghi, J., Chaganti, S. R., Johnson, T. B., & Heath, D. D. (2023). Host species and habitat shape fish-associated bacterial communities: Phylosymbiosis between fish and their microbiome. Microbiome, 11(1), 258. 10.1186/s40168-023-01697-6

93. Saghai-Maroof, M. A., Soliman, K. M., Jorgensen, R. A., & Allard, R. W. (1984). Ribosomal DNA spacer-length polymorphisms in barley: Mendelian inheritance, chromosomal location, and population dynamics. Proceedings of the National Academy of Sciences, 81(24), 8014–8018. 10.1073/pnas.81.24.8014

94. Salonen, I. S., Chronopoulou, P.-M., Nomaki, H., Langlet, D., Tsuchiya, M., & Koho, K. A. (2021). 16S rRNA Gene Metabarcoding Indicates Species-Characteristic Microbiomes in Deep-Sea Benthic Foraminifera. Frontiers in Microbiology, 12, 694406. 10.3389/fmicb.2021.694406

95. Schindelin, J., Arganda-Carreras, I., Frise, E., Kaynig, V., Longair, M., Pietzsch, T., Preibisch, S., Rueden, C., Saalfeld, S., Schmid, B., Tinevez, J.-Y., White, D. J., Hartenstein, V., Eliceiri, K., Tomancak, P., & Cardona, A. (2012). Fiji: An open-source platform for biological-image analysis. Nature Methods, 9(7), 676–682. 10.1038/nmeth.2019

96. Schuh, N. W., Carrier, T. J., Schrankel, C. S., Reitzel, A. M., Heyland, A., & Rast, J. P. (2020). Bacterial Exposure Mediates Developmental Plasticity and Resistance to Lethal Vibrio lentus Infection in Purple Sea Urchin (Strongylocentrotus purpuratus) Larvae. Frontiers in Immunology, 10, 3014. 10.3389/fimmu.2019.03014

97. Simon, J.-C., Marchesi, J. R., Mougel, C., & Selosse, M.-A. (2019). Host-microbiota interactions: From holobiont theory to analysis. Microbiome, 7(1), 5. 10.1186/s40168-019-0619-4

98. Stouthamer, R., Breeuwer, J. A. J., & Hurst, G. D. D. (1999). *Wolbachia Pipientis*: Microbial Manipulator of Arthropod Reproduction. Annual Review of Microbiology, 53(1), 71–102. 10.1146/annurev.micro.53.1.71

99. Tan, A. L., Hendricks, S. F., Carter, E. E., Buckley, K. M., & Strader, M. E. (2025). Microbial communities experienced during early development shape the host immune system and epigenome. iScience, 28(11), 113767. 10.1016/j.isci.2025.113767

100. Taylor, M., & Vega, N. M. (2021). Host Immunity Alters Community Ecology and Stability of the Microbiome in a Caenorhabditis elegans Model. mSystems, 6(2), 10.1128/msystems.00608-20. 10.1128/msystems.00608-20

101. Téfit, M. A., Budiman, T., Dupriest, A., & Yew, J. Y. (2023). Environmental microbes promote phenotypic plasticity in reproduction and sleep behaviour. Molecular Ecology, 32(18), 5186–5200. 10.1111/mec.17095

102. Wang, J., Pan, F., Soininen, J., Heino, J., & Shen, J. (2016). Nutrient enrichment modifies temperature-biodiversity relationships in large-scale field experiments. Nature Communications, 7(1), 13960. 10.1038/ncomms13960

103. Wilkins, E. M., Anderson, A. M., Buckley, K. M., & Strader, M. E. (2024). Temperature influences immune cell development and body length in purple sea urchin larvae. Marine Environmental Research, 202, 106705. 10.1016/j.marenvres.2024.106705

104. Wobbrock, J. O., Findlater, L., Gergle, D., & Higgins, J. J. (2011). The aligned rank transform for nonparametric factorial analyses using only anova procedures. Proceedings of the SIGCHI Conference on Human Factors in Computing Systems, 143–146. 10.1145/1978942.1978963

105. Wong, J. M., Kozal, L. C., Leach, T. S., Hoshijima, U., & Hofmann, G. E. (2019). Transgenerational effects in an ecological context: Conditioning of adult sea urchins to upwelling conditions alters maternal provisioning and progeny phenotype. Journal of Experimental Marine Biology and Ecology, 517, 65–77. 10.1016/j.jembe.2019.04.006

106. Yang, K., Zhang, H.-Y., Wang, P., Jin, G.-X., & Chu, D. (2024). Both symbionts and environmental factors contribute to shape the microbiota in a pest insect, Sogatella furcifera. Frontiers in Microbiology, 14, 1336345. 10.3389/fmicb.2023.1336345

107. Yao, Q., Yu, K., Liang, J., Wang, Y., Hu, B., Huang, X., Chen, B., & Qin, Z. (2019). The Composition, Diversity and Predictive Metabolic Profiles of Bacteria Associated With the Gut Digesta of Five Sea Urchins in Luhuitou Fringing Reef (Northern South China Sea). Frontiers in Microbiology, 10, 1168. 10.3389/fmicb.2019.01168

108. Yu, J., Jiang, C., Yamano, R., Koike, S., Sakai, Y., Mino, S., & Sawabe, T. (2023). Unveiling the early life core microbiome of the sea cucumber Apostichopus japonicus and the unexpected abundance of the growth-promoting Sulfitobacter. Animal Microbiome, 5(1), 54. 10.1186/s42523-023-00276-2

