## Supplemental figures for "Environmental microbial communities and host selection shape larval microbiomes"

***Supplemental figure 1.*** Alpha diversity plots

***Supplemental figure 2.*** Bar plot

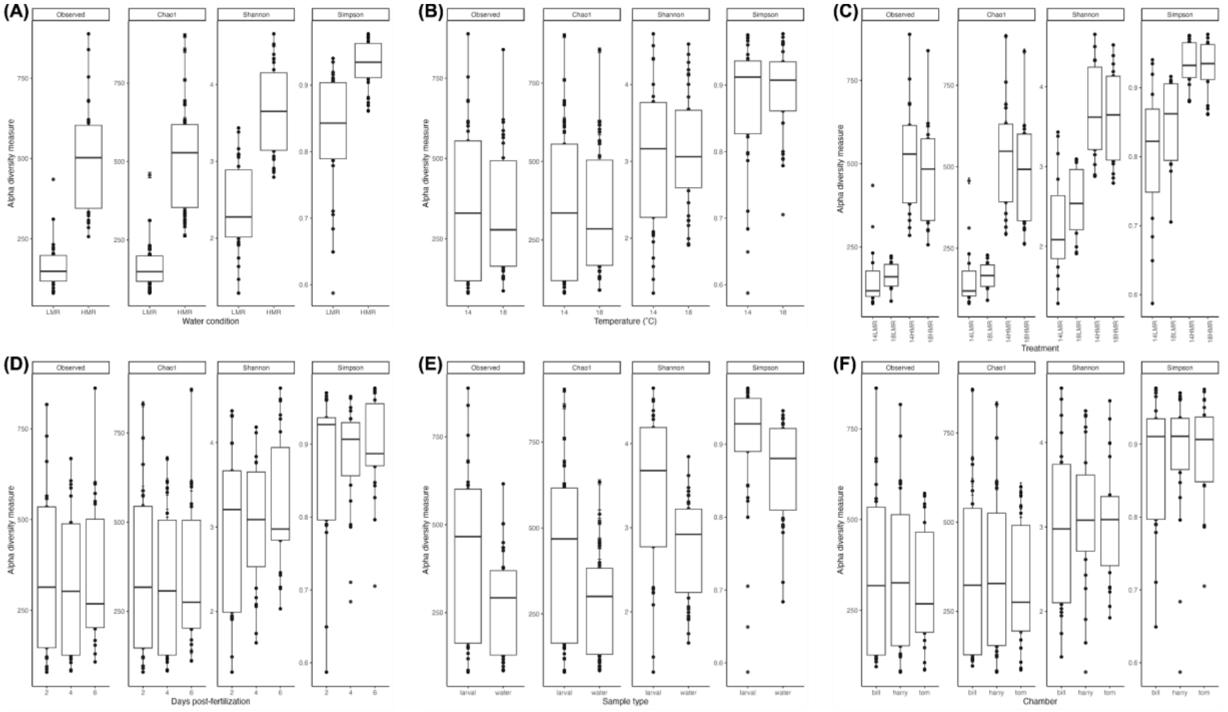


***Supplemental figure 1.*** Alpha diversity measures of Observed, Chao1, Shannon, and Simpson by water condition (A), temperature (B), treatment (C), days post-fertilization (D), sample type (E), and chamber (F).

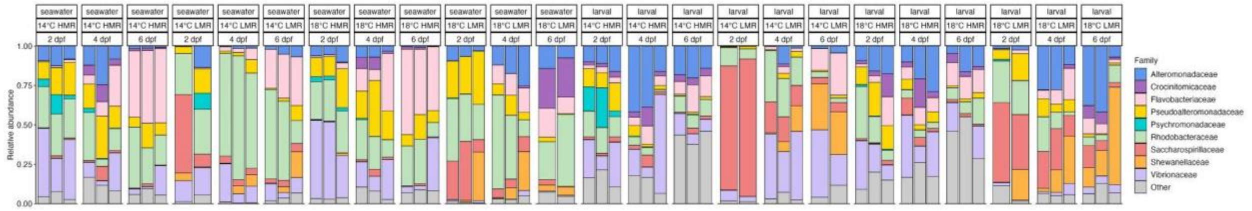


***Supplemental figure 2.*** Stacked bar plot with relative abundance of microbial communities across four treatments at all time points in seawater and larval samples. Each bar represents a sample, with colors indicating the top nine families (≥ 25% relative abundance in any single sample). "Other" consists of the remaining families grouped as one category.
